# Gene expression in patient-derived neural progenitors implicates WNT5A signaling in the etiology of schizophrenia

**DOI:** 10.1101/209197

**Authors:** Oleg V Evgrafov, Chris Armoskus, Bozena B Wrobel, Valeria N Spitsyna, Tade Souaiaia, Jennifer S. Herstein, Christopher P Walker, Joseph D Nguyen, Adrian Camarena, Jonathan R Weitz, Jae Mun ‘Hugo’ Kim, Edder Lopez Duarte, Kai Wang, George M Simpson, Janet L Sobell, Helena Medeiros, Michele T Pato, Carlos N Pato, James A Knowles

## Abstract

**BACKGROUND:** GWAS of schizophrenia demonstrated that variations in the non-coding regions are responsible for most of common variation heritability of the disease. It is hypothesized that these risk variants alter gene expression. Thus, studying alterations in gene expression in schizophrenia may provide a direct approach to understanding the etiology of the disease. In this study we use <u>C</u>ultured <u>N</u>eural progenitor cells derived from <u>O</u>lfactory <u>N</u>euroepithelium (CNON) as a genetically unaltered cellular model to elucidate the neurodevelopmental aspects of schizophrenia.

**METHODS:** We performed a gene expression study using RNA-Seq of CNON from 111 controls and 144 individuals with schizophrenia. Differentially expressed (DEX) genes were identified with DESeq2, using covariates to correct for sex, age, library batches and one surrogate variable component.

**RESULTS:** 80 genes were DEX (FDR<10%), showing enrichment in cell migration, cell adhesion, developmental process, synapse assembly, cell proliferation and related gene ontology categories. Cadherin and Wnt signaling pathways were positive in overrepresentation test, and, in addition, many genes are specifically involved in Wnt5A signaling. The DEX genes were significantly, enriched in the genes overlapping SNPs with genome-wide significant association from the PGC GWAS of schizophrenia (PGC SCZ2). We also found substantial overlap with genes associated with other psychiatric disorders or brain development, enrichment in the same GO categories as genes with mutations *de novo* in schizophrenia, and studies of iPSC-derived neural progenitor cells.

**CONCLUSIONS:** CNON cells are a good model of the neurodevelopmental aspects of schizophrenia and can be used to elucidate the etiology of the disorder.

## Introduction

Schizophrenia (SCZ) is a devastating psychiatric disorder with an estimated lifetime prevalence of 0.55% worldwide (1). With heritability of about 81% (2), genetics plays a critical role in the disease’s etiology, however the mechanism by which genetic variation contributes to the disease is unknown. Many direct and indirect pieces of evidence indicate that aberrations in brain development are major contributors (3–5). Post-mortem studies of the brains of adults with SCZ provide important information about functional changes associated with the disease, but it is highly unlikely that the gene expression profiles of differentiated cells can provide a full picture of changes in neurodevelopment. Different biological models, such as hiPSC-derived neural progenitor cells (6) and olfactory epithelium derived cells/tissue (7,8) have been suggested to study neurodevelopmental aberrations in schizophrenia. For this study, we have used <u>C</u>ultured <u>N</u>eural progenitor cells derived from <u>O</u>lfactory <u>N</u>euroepithelium (CNON) of individuals with, and without, schizophrenia (9). These cells are not genetically modified neural progenitors, actively divide, and migrate in 2D and 3D cultures. Here we present a study of transcriptome expression profiles using RNA-Seq (strand-specific, rRNA-depleted total RNA) in CNON lines derived from 144 SCZ and 111 control (CTL) individuals.

## Materials and Methods

### Recruitment and sample collection

This study was approved by the University of Southern California and SUNY Downstate IRBs. We collected samples from 144 patients with DSM-IV criteria for schizophrenia (SCZ) and 111 controls. Most patients and control subjects were recruited from participants of the Genomic Psychiatry Cohort (GPC) study (1R01MH085548) (10), and a few patients were recruited through Los Angeles county/University of Southern California outpatient psychiatric clinic. Given the common variation overlap of SCZ and Major Depressive Disorder (MDD) (11), we excluded controls that endorsed either MDD probe question on the GPC screener (10), effectively removing individuals with a potential history of MDD.

### Cell culture and transcriptome sequencing

We developed cell cultures from olfactory biopsies (12) as described previously (9).

In brief, each biopsy sample was dissected into pieces approximately 1mm3 in size, placed on the surface of a 60mm tissue culture dish coated by Matrigel basement membrane (MBM) (BD Bioscience, San Jose, California, USA) reconstituted in Coon’s medium (1:2), and then every piece of tissue sample was covered by a droplet of full-strength MBM. After MBM gelatinizes, 5 ml of medium 4506 (Ghanbari et al., 2004) was added. Within 1–4 weeks of culturing, CNON cells were observed to grow out of the embedded pieces of tissue. Due to unique ability to grow through Matrigel, neural progenitors often populate large areas without presence of other cell types (supplemental Figure S1). Outgrown cells with a neuronal phenotype were then physically isolated using cloning cylinders and dislodged using Dispase (BD Bioscience). Cells collected from inside the cloning cylinders were further grown on tissue culture Petri dishes covered by reconstituted MDM in 4506 medium. RNA was purified from ∼400,000 cells grown on 6 cm Petri dishes (∼90% of confluence), using the Direct-Zol RNA MiniPrep kit (Zymo Research), according to manufacturer’s protocol. RNA libraries were prepared in batches of 24-48 samples with approximately equal numbers of cases and controls with TruSeq Stranded Total RNA LT Library preparation kits with Ribo-Zero Gold (Illumina) according to manufacturer’s protocol, using a Hamilton STARlet liquid handling robot to increase library preparation consistency. Equimolar pools of at least 4 libraries, containing both cases and controls, were constructed after quantification using the KAPA Library Quantification Kit (Kapa Biosystems), and sequenced together using HiSeq2000 DNA Sequencers (Illumina) with 100 bp single-end reads. On average, each sample was run in 3.93 lanes across 3.91 flow cells, to reduce potential channel and flow cell bias.

### Mapping and assignment of reads to genes

We performed pre-mapping QC, read trimming, mapping and assignment to the sense strand of gene models (GenCode v22, max mismatch=6) using GT-FAR v12 (https://genomics.isi.edu/gtfar) (see Supplemental Materials for detail). The number of uniquely mapped reads uniquely assigned to each gene model was used as a proxy of gene expression in DEX gene analysis.

### RT-qPCR

Reverse transcription quantitative Polymerase Chain Reaction (RT-qPCR) was performed in duplicates using the Biomark HD (Fluidigm) on a Flex Six Gene Expression IFC (Integrated Fluidic Circuit), according to manufacturer’s protocols and using the recommended reagents. Normalized relative expression (to *ACTB*) was calculated using the ΔΔC_t_ method (13), and log-transformed expression values were analyzed by ANOVA controlling for sex.

### Differential gene expression analysis

We performed main differential gene expression (DEX) analysis between SCZ and controls using DESeq2 v1.16.1 (14) in R v3.4.1, an algorithm assessing difference between mean gene expression in groups using a generalized linear model and assuming a negative binomial distribution of RNA-Seq reads. The analysis used covariates of sex, age, 4 library batches and 1 surrogate variable (SVA v3.24.4). Procedures used to arrive at this covariate set are further described in Supplemental Materials. DESeq2’s built-in normalization process was used.

The analysis was done on 23,920 expressed genes, defined as having on average at least 3.5 reads per sample, based on the density plot of log-transformed baseMean for all genes (Supplemental Figure S2). Resulting p-values were adjusted for multiple comparisons based on the Benjamini-Hochberg False Discovery Rate (FDR) using the p.adjust function in R. Transcripts Per Million transcripts (TPM) values were calculated by dividing the mean number of unnormalized reads uniquely mapped to a gene by the median transcript length (15) in Gencode 22 for that gene and normalizing the resulting values so they sum to one million (16). Hierarchical clustering analysis was done in R using the ‘hclust’ function where the distance is calculated as one minus the Pearson correlation coefficient and using average linkage.

### Permutation analysis

To test that our findings are not due to random variation we performed two forms of permutation analysis. First, 1,000 analyses were run under the same conditions as the main analysis, except with labels for diagnosis randomly permuted for all samples. Second, 250 analyses each of random subsets in 4 different configurations were analyzed: random half of cases vs. other half of cases, and random half of controls vs. other half of controls (null comparisons); and, due to the difference in number of cases and controls, random half of cases vs. random same number of controls, and random half of controls vs. random same number of cases (case/control comparisons). The numbers of differentially expressed (DEX) genes in the permuted analyses were compared to the number seen in the original, unpermuted analysis by Wilcoxon test. The number of DEX genes in the null comparisons was compared to the number DEX in the case/control comparisons by Mann-Whitney test.

### GWAS enrichment analysis

We identified GWAS variants’ p-values within a given gene based on the PGC SCZ2 dataset (https://www.med.unc.edu/pgc/results-and-downloads) and Gencode 22 annotation back-ported to human reference genome *GRCh37* (hg19) to match coordinates used in GWAS. Fisher’s Exact Test was used to test for enrichment of genes co-localized with genome-wide significant (p<5×10^−8^) GWAS peaks and calculate estimated odds-ratio. Due to a very broad peak in the HLA region (25 - 34 Mb of chromosome 6 in *GRCh37* coordinates), genes from this region were removed from the analysis, as were genes from the Y chromosome.

### WGCNA

Weighted Gene Correlation Network Analysis (WGCNA) (17) was performed for all expressed genes on residuals after correction for effects of three explicit batches as well as one surrogate variable, with a soft-threshold power of 5 to achieve approximate scale-free topology (SFT R^2^>0.85; truncated R^2^>0.95), and a minimum module size of 50 genes. Modules of co-expressed genes produced by the algorithm were tested for enrichment of DEX genes by Fisher’s exact test and for correlation of the eigengene with diagnosis (SCZ vs. control), after controlling for sex and age (linear model, no interactions); the p-value cutoff was adjusted for multiple comparisons. Gene set enrichment analysis was applied to modules that had a significant p-value on the aforementioned statistical tests.

## Results

We collected samples of olfactory neuroepithelium and established CNON cell lines from 144 individuals with DSM-IV SCZ and 111 CTL (Supplemental Table S1) (9,12). Strand-specific RNA-Seq of total RNA was performed, resulting in an average of 23.38 million (7.1 – 106.7 million) uniquely mapped reads per sample, after exclusion of rRNA and mitochondrial genes. DESeq2 was used to normalize the read counts assigned to expressed genes and perform differential gene expression analysis.

### Characterization of CNON cells

The RNA-Seq data in this study are consistent with our previous observations using Affymetrix Human Exon 1.0 ST arrays (9) that CNON lines are neural progenitors (Supplemental table S2). In order to determine the period of human brain development the CNON lines most resemble, we compared RNA-Seq data from CNON to 647 poly-A RNA-Seq 76 bp datasets from post-mortem human brain samples across 41 individuals, 26 brain regions and 10 developmental stages (from “Early Fetal” to “Middle Adulthood”) (www.brainspan.org, (18)). To mitigate the differences in RNA-Seq methodology, we transformed the gene expression values of each CNON sample using the coordinates described by the principal components of the BrainSpan data. The first principle component of the BrainSpan data roughly corresponds to developmental age and separates the pre- and post-natal samples (Supplemental Figure S3). The CNON samples form a tight cluster within the prenatal samples, particularly those from the mid fetal period (weeks 13-24, second trimester).

### Genes differentially expressed in neural progenitor cells in SCZ

Using our analysis model (see Supplemental Materials for details), 80 genes were DEX between SCZ and CTL at a false discovery rate (FDR) of 10%, corresponding to a maximum p-value of 3.3×10^−4^ (Table 1; complete gene list in Supplemental Table S3). The average fold change was 1.8 (range 1.08 to 9.09) (Figure 1) at expression levels of 0.07 to 552 TPM.

**Table 1.** Genes significantly DEX between SCZ and Control at FDR < 0.1. To facilitate the comparison of expression of each gene, normalized read counts were transformed to transcripts per million transcripts (TPM) (16), using the median transcript length (15) in Gencode release 22 for each gene as gene length. Gene symbols and gene types are taken from Gencode release 27 where available. Notes: 1) Gene lies under genome-wide significant PGC SCZ2 GWAS peak (19); 2) *De novo* non-silent mutations in the gene has been identified in patients with SCZ; 3) *De novo* missense mutation(s) in the gene has been identified in patients with autism spectrum disorder; 4) CNVs were identified in multiple patients with SCZ, 5) gene found significantly associated with psychiatric disorder or related traits (other than PGC SCZ2 GWAS).

| Ensembl ID | Gene Symbol | Gene Type | TPM | Log2 Fold-Change | FDR | Notes |
| --- | --- | --- | --- | --- | --- | --- |
| ENSG00000278099 | *U1* | snRNA | 4.84 | 3.184 | $9.77 \times 10^{-10}$ | |
| ENSG00000170627 | *GTSF1* | protein_coding | 0.362 | 2.162 | $5.50 \times 10^{-07}$ | |
| ENSG00000149564 | *ESAM* | protein_coding | 0.347 | -1.423 | $3.79 \times 10^{-06}$ | 1, 2, 5 |
| ENSG00000115461 | *IGFBP5* | protein_coding | 552 | 1.045 | $5.54 \times 10^{-04}$ | |
| ENSG00000124749 | *COL21A1* | protein_coding | 0.926 | 1.184 | $2.31 \times 10^{-03}$ | |
| ENSG00000180447 | *GAS1* | protein_coding | 24.3 | 0.842 | $2.47 \times 10^{-03}$ | |
| ENSG00000157766 | *ACAN* | protein_coding | 0.217 | -1.232 | $5.69 \times 10^{-03}$ | |
| ENSG00000164532 | *TBX20* | protein_coding | 0.115 | -1.509 | 5.77x10<sup>-03</sup> |  |
| ENSG00000135914 | *HTR2B* | protein_coding | 2.74 | 1.169 | 5.77x10<sup>-03</sup> |  |
| ENSG00000008517 | *IL32* | protein_coding | 2.33 | 1.044 | 5.77x10<sup>-03</sup> |  |
| ENSG00000165197 | *VEGFD* | protein_coding | 0.614 | 1.125 | 6.66x10<sup>-03</sup> |  |
| ENSG00000141469 | *SLC14A1* | protein_coding | 7.48 | -1.679 | 6.66x10<sup>-03</sup> |  |
| ENSG00000260604 | *AL590004.4* | lincRNA | 0.602 | 0.813 | 9.83x10<sup>-03</sup> |  |
| ENSG00000114251 | *WNT5A* | protein_coding | 186 | 0.521 | 9.83x10<sup>-03</sup> |  |
| ENSG00000184731 | *FAM110C* | protein_coding | 0.591 | -1.003 | 2.71x10<sup>-02</sup> | 5 |
| ENSG00000162434 | *JAK1* | protein_coding | 81.7 | 0.180 | 2.98x10<sup>-02</sup> | 3 |
| ENSG00000143631 | *FLG* | protein_coding | 0.127 | -1.248 | 3.41x10<sup>-02</sup> | 3 |
| ENSG00000133048 | *CHI3L1* | protein_coding | 1.036 | 1.513 | 3.49x10<sup>-02</sup> |  |
| ENSG00000118689 | *FOXO3* | protein_coding | 9.45 | 0.207 | 3.49x10<sup>-02</sup> | 1, 5 |
| ENSG00000226237 | *GAS1RR* | lincRNA | 0.719 | 0.503 | 3.49x10<sup>-02</sup> |  |
| ENSG00000172156 | *CCL11* | protein_coding | 1.45 | 1.426 | 3.67x10<sup>-02</sup> |  |
| ENSG00000230552 | *AC092162.2* | lincRNA | 0.460 | 0.707 | 4.12x10<sup>-02</sup> |  |
| ENSG00000203648 | *AC007618.1* | processed_pseudogene | 0.662 | 0.522 | 4.53x10<sup>-02</sup> |  |
| ENSG00000076706 | *MCAM* | protein_coding | 15.52 | -1.012 | 4.53x10<sup>-02</sup> | 5, 3 |
| ENSG00000109819 | *PPARGC1A* | protein_coding | 2.09 | 0.715 | 4.53x10<sup>-02</sup> |  |
| ENSG00000188227 | *ZNF793* | protein_coding | 8.96 | 0.244 | 4.61x10<sup>-02</sup> |  |
| ENSG00000145819 | *ARHGAP26* | protein_coding | 16.85 | 0.324 | 4.61x10<sup>-02</sup> | 3 |
| ENSG00000120324 | *PCDHB10* | protein_coding | 0.096 | 0.914 | 4.64x10<sup>-02</sup> |  |
| ENSG00000135250 | *SRPK2* | protein_coding | 63.78 | 0.107 | 4.91x10<sup>-02</sup> | 1, 5 |
| ENSG00000233476 | *EEF1A1P6* | processed_pseudogene | 0.550 | 0.382 | 4.91x10<sup>-02</sup> |  |
| ENSG00000112852 | *PCDHB2* | protein_coding | 1.22 | 0.647 | 4.91x10<sup>-02</sup> |  |
| ENSG00000189058 | *APOD* | protein_coding | 3.91 | 0.885 | 4.91x10<sup>-02</sup> |  |
| ENSG00000120337 | *TNFSF18* | protein_coding | 3.05 | 1.216 | 4.91x10<sup>-02</sup> |  |
| ENSG00000197181 | *PIWIL2* | protein_coding | 0.520 | -0.718 | 4.91x10<sup>-02</sup> |  |
| ENSG00000279118 | *AC093535.2* | TEC | 7.38 | 0.454 | 4.91x10<sup>-02</sup> |  |
| ENSG00000277232 | *GTSE1-AS1* | lincRNA | 0.858 | -0.423 | 4.91x10<sup>-02</sup> |  |
| ENSG00000235531 | *MSC-AS1* | antisense_RNA | 36.55 | 0.343 | 5.50x10<sup>-02</sup> |  |
| ENSG00000170921 | *TANC2* | protein_coding | 19.16 | 0.213 | 5.71x10<sup>-02</sup> | 2, 5 |
| ENSG00000154721 | *JAM2* | protein_coding | 7.324 | 0.712 | 5.92x10<sup>-02</sup> |  |
| ENSG00000137558 | *PI15* | protein_coding | 0.147 | 1.091 | 5.92x10<sup>-02</sup> | 2 |
| ENSG00000124107 | *SLPI* | protein_coding | 1.241 | -2.265 | 5.92x10<sup>-02</sup> |  |
| ENSG00000109158 | *GABRA4* | protein_coding | 0.362 | 0.483 | 5.92x10<sup>-02</sup> |  |
| ENSG00000154229 | *PRKCA* | protein_coding | 89.5 | 0.191 | 5.92x10<sup>-02</sup> | 3 |
| ENSG00000113205 | *PCDHB3* | protein_coding | 0.136 | 0.846 | 5.92x10<sup>-02</sup> | 5 |
| ENSG00000255583 | *AC084357.3* | transcribed_unprocessed_pseudogene | 0.819 | -0.824 | 5.92x10<sup>-02</sup> |  |
| ENSG00000183690 | *EFHC2* | protein_coding | 0.162 | 0.908 | $5.92 \times 10^{-02}$ | |
| ENSG00000176896 | *TCEANC* | protein_coding | 1.374 | 0.226 | $5.92 \times 10^{-02}$ | |
| ENSG00000021645 | *NRXN3* | protein_coding | 0.615 | -0.797 | $6.20 \times 10^{-02}$ | |
| ENSG00000258071 | *ARL2BPP2* | processed_pseudogene | 2.53 | 0.300 | $6.42 \times 10^{-02}$ | |
| ENSG00000104903 | *LYL1* | protein_coding | 0.485 | -0.502 | $6.93 \times 10^{-02}$ | 5 |
| ENSG00000169122 | *FAM110B* | protein_coding | 17.59 | 0.387 | $7.28 \times 10^{-02}$ | |
| ENSG00000245680 | *ZNF585B* | protein_coding | 30.62 | 0.194 | $7.28 \times 10^{-02}$ | |
| ENSG00000169306 | *IL1RAPL1* | protein_coding | 0.074 | -0.899 | $7.28 \times 10^{-02}$ | 3, 4 |
| ENSG00000068305 | *MEF2A* | protein_coding | 24.27 | 0.212 | $7.81 \times 10^{-02}$ | |
| ENSG00000234155 | *LINC02535* | lincRNA | 0.797 | -0.376 | $7.93 \times 10^{-02}$ | |
| ENSG00000272674 | *PCDHB16* | protein_coding | 0.648 | 0.610 | $7.93 \times 10^{-02}$ | 3 |
| ENSG00000111371 | *SLC38A1* | protein_coding | 137 | 0.273 | $7.93 \times 10^{-02}$ | |
| ENSG00000236535 | *RC3H1-IT1* | sense_intronic | 2.202 | 0.334 | $7.93 \times 10^{-02}$ | |
| ENSG00000164265 | *SCGB3A2* | protein_coding | 5.455 | 0.395 | $7.93 \times 10^{-02}$ | |
| ENSG00000163815 | *CLEC3B* | protein_coding | 6.303 | 0.691 | $7.93 \times 10^{-02}$ | |
| ENSG00000223361 | *FTH1P10* | transcribed_processed_pseudogene | 0.732 | -0.906 | $8.33 \times 10^{-02}$ | |
| ENSG00000139508 | *SLC46A3* | protein_coding | 4.206 | 0.302 | $8.33 \times 10^{-02}$ | |
| ENSG00000173947 | *PIFO* | protein_coding | 0.142 | 0.517 | $8.52 \times 10^{-02}$ | |
| ENSG00000146453 | *PNLDC1* | protein_coding | 0.181 | 1.274 | $8.52 \times 10^{-02}$ | |
| ENSG00000182240 | *BACE2* | protein_coding | 32.35 | 0.304 | $8.58 \times 10^{-02}$ | |
| ENSG00000113269 | *RNF130* | protein_coding | 24.13 | 0.173 | $8.78 \times 10^{-02}$ | |
| ENSG00000139926 | *FRMD6* | protein_coding | 284 | 0.263 | $8.81 \times 10^{-02}$ | |
| ENSG00000260522 | *AC106785.1* | processed_pseudogene | 2.283 | 0.385 | $8.83 \times 10^{-02}$ | |
| ENSG00000279250 | *AC022919.1* | TEC | 1.262 | 0.419 | $9.29 \times 10^{-02}$ | |
| ENSG00000226261 | *AC064836.1* | processed_pseudogene | 3.389 | -0.519 | $9.88 \times 10^{-02}$ | |
| ENSG00000189067 | *LITAF* | protein_coding | 452 | 0.250 | $9.88 \times 10^{-02}$ | |
| ENSG00000164244 | *PRRC1* | protein_coding | 34.65 | 0.131 | $9.88 \times 10^{-02}$ | |
| ENSG00000272769 | *AC097532.2* | lincRNA | 0.237 | 0.476 | $9.88 \times 10^{-02}$ | |
| ENSG00000174697 | *LEP* | protein_coding | 1.893 | 1.342 | $9.88 \times 10^{-02}$ | |
| ENSG00000157216 | *SSBP3* | protein_coding | 4.987 | 0.264 | $9.88 \times 10^{-02}$ | 2 |
| ENSG00000261824 | *LINC00662* | lincRNA | 21.58 | 0.205 | $9.88 \times 10^{-02}$ | |
| ENSG00000106459 | *NRF1* | protein_coding | 8.061 | -0.142 | $9.88 \times 10^{-02}$ | |
| ENSG00000162383 | *SLC1A7* | protein_coding | 0.142 | 0.828 | $9.88 \times 10^{-02}$ | |
| ENSG00000228495 | *LINC01013* | lincRNA | 1.26 | 0.921 | $9.90 \times 10^{-02}$ | |
| ENSG00000197928 | *ZNF677* | protein_coding | 20.20 | 0.171 | $9.92 \times 10^{-02}$ | |

**Figure 1.**
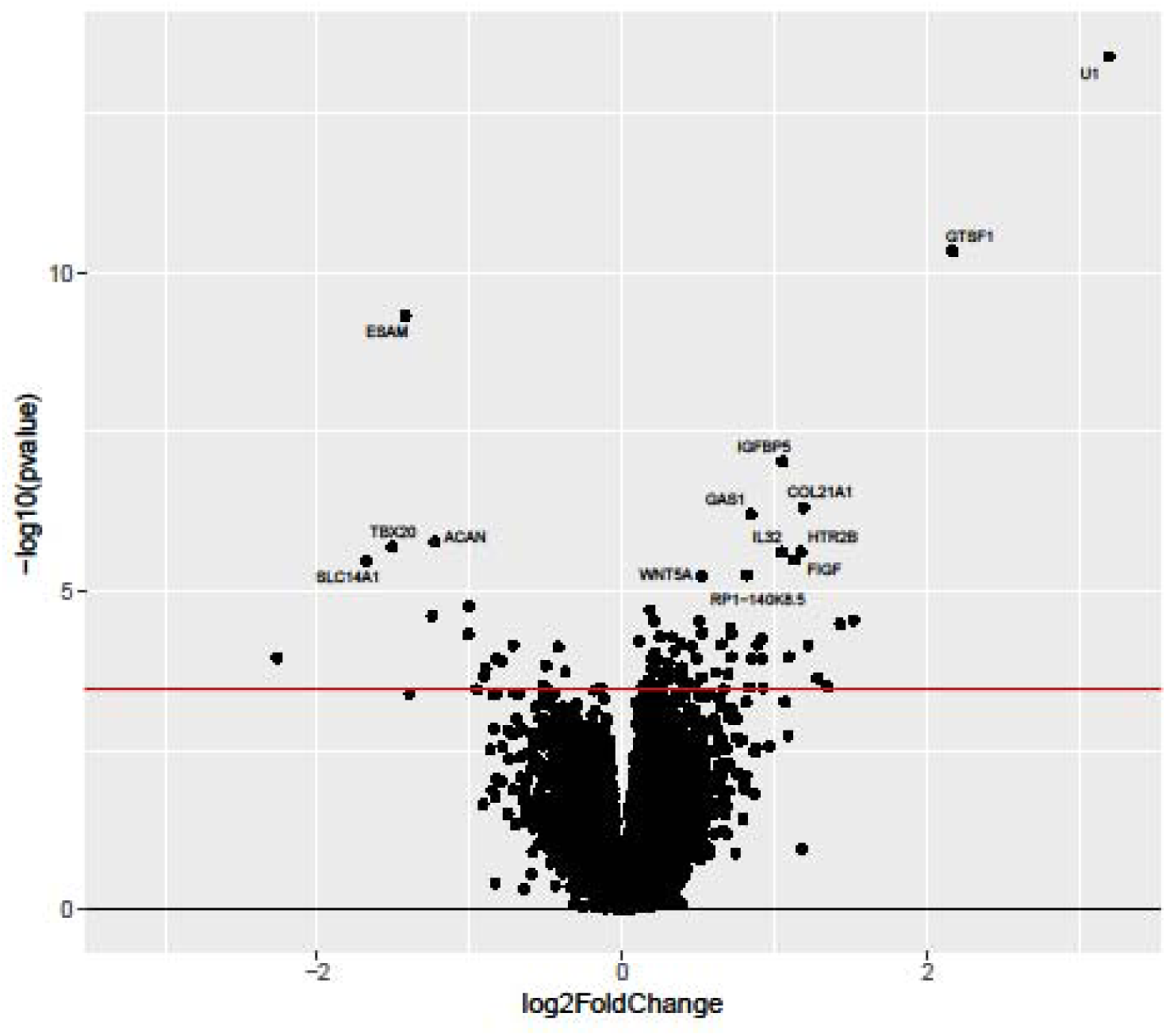
Volcano plot for SCZ vs. CTL DEX comparison. Genes with raw p < 10^−5^ are labeled. Red line indicates FDR < 10%.

To evaluate the accuracy of RNA-Seq gene quantification and DEX analysis, we compared results with RT-qPCR. For that purpose, we used a subset of 146 samples and performed DEX analysis on RNA-Seq data on only these samples. From this list of DEX genes we selected 5 genes (*CCL8, HTR2B, PLAT, PPARGC1A*, and *VAV3*) to include genes with fold change differences in both directions and spanning a range of gene expression levels. Expression of these genes and *ACTB* (used for normalization) was assessed by RT-qPCR on the same set of 146 samples. Expression data from RT-qPCR and RNA-Seq were highly correlated within each gene (mean r=0.75, range 0.63 - 0.90, all p-values <2×10^−15^) (Supplemental Figure S4A-E), and mean expression of all six genes had correlation r=0.943 (p=0.0047) (Supplemental Figure S4F). DEX of 4 of 5 genes was replicated, while one gene (*PLAT*) did not reach significance (p=0.15) but shows a trend in the same direction as that seen in RNA-Seq (Supplemental Figure S4GH).

### Permutation analysis of differential expression

To assess the probability that DEX findings could be due to random statistical variation, we performed two forms of permutation analysis. In the first, we randomly permuted the diagnosis labels, but held all other factors constant. We found a median of 11 differentially expressed genes at FDR<10%, which is significantly lower (p<2×10^−16^, Wilcoxon signed rank test) than in the unpermuted data. In the second permutation analysis, where we permute subsets of the data to compare case-control comparisons to null comparisons (SCZ vs. SCZ and CTL vs. CTL) (see Methods for detail), the null comparisons resulted in significantly fewer differentially expressed genes at FDR<10% (median=4, mean=38.45), as compared to case/control comparisons (median=9, mean=152.98); p<3.6×10^−15^ by Mann-Whitney test (Supplemental Figure S5 and Supplemental Materials). These permutation analyses strongly suggest that the majority of genes found to be DEX are not likely to be false positives.

### GWAS enrichment analysis

It is thought that causal variants in SCZ GWAS loci regulate expression of genes involved in the etiology of SCZ (20,21). Previous studies have shown that regulatory variants are likely to be localized near to or within the genes they directly regulate (22), we tested the hypothesis that DEX genes are more likely to be co-localized with genome-wide significant (p<5×10^−8^) variants from the PGC SCZ2 GWAS (19), excluding the HLA region of chromosome 6. Three DEX genes (*ESAM, FOXO3*, and *SRPK2*) were found to overlap independent genome-wide significant variants (Fisher’s exact test, odds-ratio=3.8, p=0.049) (Supplemental Figure S6). Two other genes just missed being overlapped with genome-wide significant peaks. DEX gene *IL1RAPL1* is almost genome-wide significant (p=5.3×10^−8^), and *AC007618.3* sits within an intron of *CACNA1C*, in which another intron contains one of the most significant GWAS signals (19). For comparison, genes that were DEX (FDR<10%) based on either sex or age were not found to be enriched for overlap with SCZ GWAS variants (p>0.05).

The index SNP of another GWAS-significant locus (rs56972983, chromosome 5) is an eQTL for *PCDHB16*, in GTEx v7 (aorta). Three other DEX protocadherin genes, *PCDHB10, PCDHB2* and *PCDHB3*, also have eQTLs within a broader region of this GWAS peak. It is noteworthy that all 48 protocadherin genes transcribed in CNON from the alpha, beta and gamma clusters, are expressed at a higher level in SCZ.

### Gene set enrichment, pathway and network analyses of DEX genes

Gene set enrichment analysis (GSEA) shows significant (corrected p<0.05) enrichment of the DEX genes in 47 GO terms from 10 categories (related groups of terms) (Table 2, Supplemental Table S4), including cell migration, cell adhesion, developmental process, cell proliferation, synapse assembly and PANTHER pathway analysis (23) shows over-representation of DEX genes involved in the cadherin and Wnt pathways (FDR=1.39% and 1.91% respectively). Hierarchical clustering of DEX genes further supports the finding of involvement of Wnt signaling (Figure 2). The most prominent cluster of correlated genes includes: *WNT5A*, the most expressed Wnt ligand gene in CNON; Leptin and *JAK1*, which regulate WNT5A expression; *MCAM*, which encodes a strong WNT5A receptor; *PRKCA*, which encodes protein kinase C, a protein involved in non-canonical Wnt/calcium pathways; *PPARGC1A* and *FOXO3*, which interact with β-catenin, a key molecule in Wnt signaling (Figure 3). All together these results strongly indicate alteration of WNT5A signaling as one of potential causes of SCZ.

**Table 2:** Gene Ontology enrichment results based on 80 DEX genes (FDR < 10%). GSEA done by g:Profiler (24) using the ordered query option that takes into account which genes are more significant. Hierarchical filtering was applied to produce only the most significant term per parent term. Complete table of significant terms is presented in Supplemental Table S4. P-values were corrected for multiple comparisons by algorithm g:SCS, the default option in g:Profiler.

| GO Term Name | GO Term ID | Corrected p-value | DEX Genes |
| --- | --- | --- | --- |
| Angiogenesis | GO:0001525 | 3.53E-06 | 11 genes |
| Cell migration | GO:0016477 | 9.48E-06 | 15 genes |
| Cell adhesion | GO:0007155 | 0.000125 | 16 genes |
| Synapse assembly | GO:0007416 | 0.00031 | 7 genes: *IL1RAPL1*, *NRXN3*, *PCDHB2*, *PCDHB3*, *PCDHB10*, *PCDHB16*, *WNT5A* |
| Positive regulation of endothelial cell proliferation | GO:0001938 | 0.000864 | 5 genes: *CCL11*, *HTR2B*, *PRKCA*, *VEGFD*, *WNT5A* |
| Calcium-dependent cell-cell adhesion via plasma membrane cell adhesion molecules | GO:0016339 | 0.00283 | 4 genes: *PCDHB2*, *PCDHB3*, *PCDHB10*, *PCDHB16* |
| Regulation of cell proliferation | GO:0042127 | 0.00291 | 12 genes |
| Striated muscle hypertrophy | GO:0014897 | 0.0094 | 5 genes: *HTR2B*, *IGFBP5*, *LEP*, *MEF2A*, *PRKCA* |
| Regulation of lipoprotein lipid oxidation | GO:0060587 | 0.0425 | 2 genes: *APOD*, *LEP* |
| Mammary gland morphogenesis | GO:0060443 | 0.05 | 3 genes: *CCL11*, *IGFBP5*, *WNT5A* |

**Figure 2.**
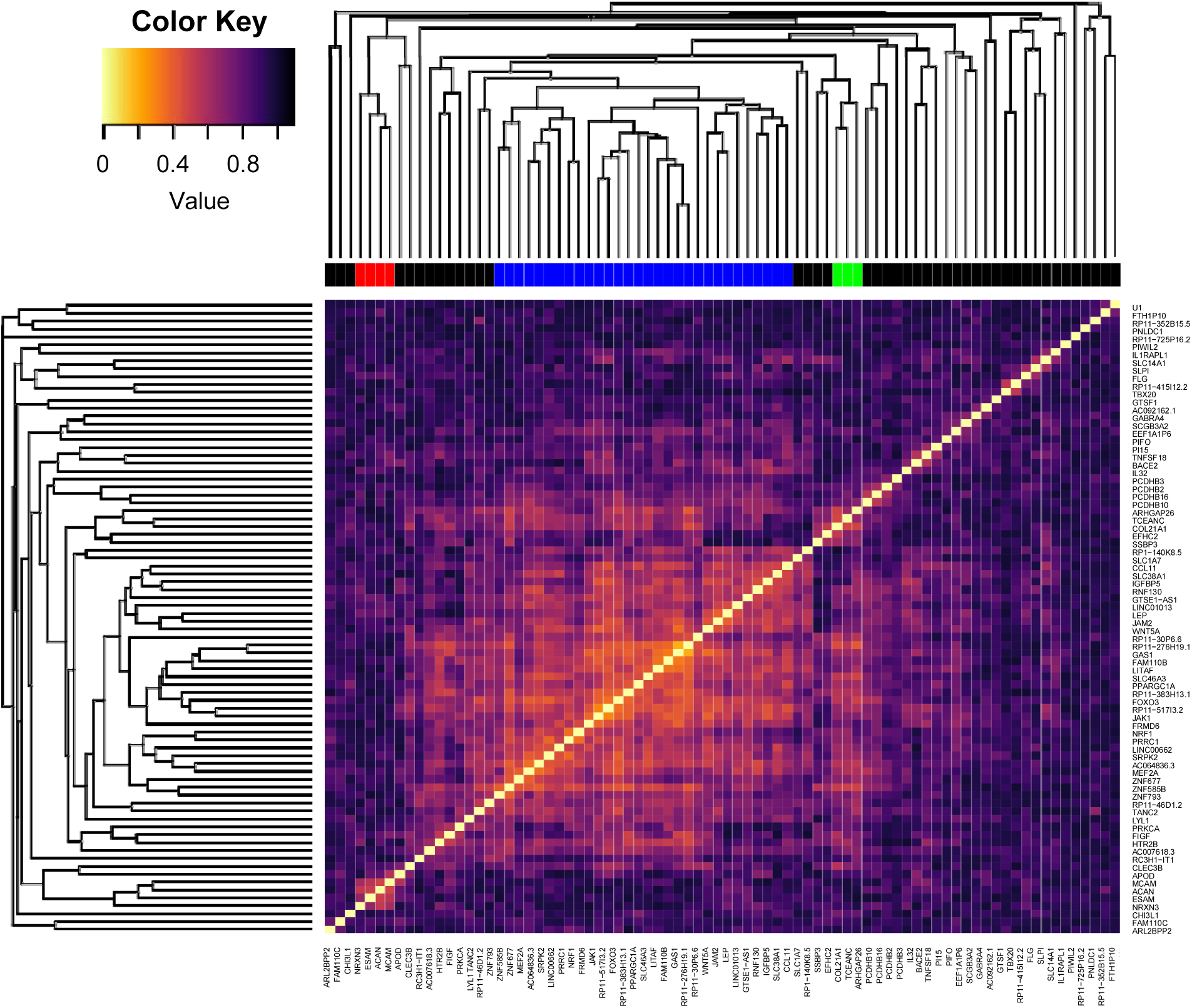
Heatmap and hierarchical clustering of DEX genes. Clustering was performed using average linkage and a distance of one minus the absolute value of the Pearson correlation coefficient. Lighter color indicates higher correlation. Genes were assigned to a group (colors in bar above heatmap) based on a cutoff at clustering distance of 0.6.

**Figure 3.**
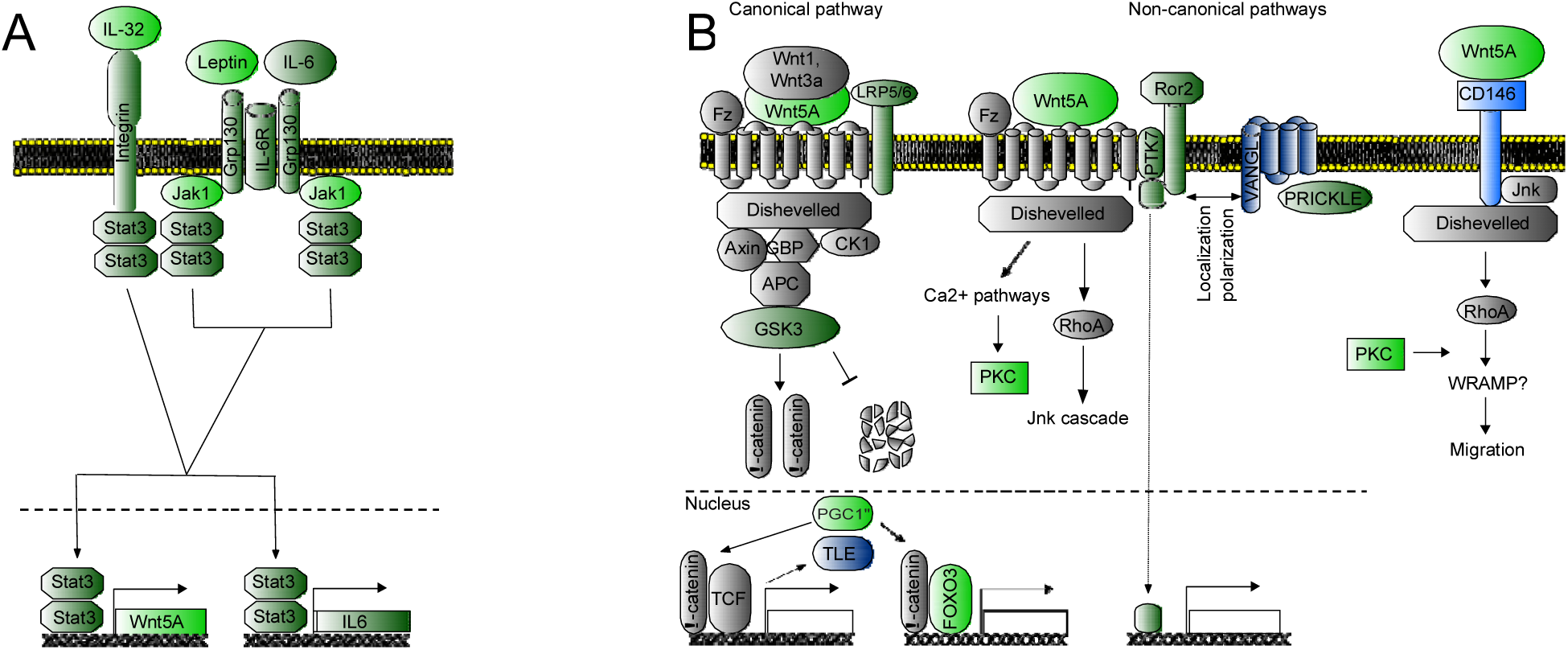
Wnt signaling branches involving genes with differential expression between SCZ and control groups. Light green: DEX (FDR<10%), over-expressed in SCZ; light blue: DEX (FDR<10%), over-expressed in SCZ; dark-green: over-expressed in SCZ, p<0.05 (nominally significant); dark-blue: under-expressed in SCZ, p<0.05; grey: not differentially expressed (p>0.05); empty box: unknown genes. **(A) Regulation of WNT5A expression.** *WNT5A* is regulated by JAK1, LEP, STAT3, IL6ST, and IL6, which are known as the STAT3-WNT5A signaling loop (25). All genes corresponding to these proteins have higher expression in SCZ group (p<0.05), the first two genes being DEX, pointing to specific mechanism of elevated *WNT5A* expression in SCZ. *IL32*, another DEX gene, may regulate *IL6* production (26) and/or directly contribute to increased expression of *WNT5A* through activating *STAT3*. **(B) Wnt signaling pathways.** The information about involvement of WNT5A in induction of canonical signaling is conflicting, but it is generally agreed that it is mostly involved in non-canonical signaling. Three genes involved in canonical pathway show changes in expression in line with enhanced signaling. *LRP5*, co-receptor of fizzled, has higher expression in SCZ, while *TLE*, repressor of β-catenin target genes, has lower expression (p<0.05). *GSK3B* was previously implicated in bipolar disorder (27), another psychiatric disease which shares some genetic susceptibility with SCZ. *GSK3B* did not reach the level of transcriptome-wide significance in our study (#294 in rank of significance, p=0.002), but direction of expression change is in line with increased Wnt signaling. DEX gene *FOXO3* encodes a transcription factor, which interferes with transcription factors of canonical Wnt signaling (28), is over-expressed in SCZ and its expression highly correlates with *WNT5A* expression in CNON. PPARG coactivator 1 alpha, also encoded by a DEX gene, alters expression of transcription factors involved in canonical Wnt signaling (29). **Non-canonical WNT5A signaling pathways.** Frizzled, Dishevelled, VANGL and PRICKLE are core proteins of the Planar Cell Polarity pathway, activated by WNT5A, with ROR2 (p<0.05) and PTK7 (p<0.05) being significant co-receptors (25,30). Complexes of Frizzled and Dishevelled are localized on the opposite side of cells from VANGL and PRICKLE, polarizing cells and playing a role in polarized movement (31). In CNON VANGL proteins are presented almost exclusively by under-expressed VANGL1 (p<0.05), while both *PRICKLE1* and *PRICKLE2* genes are expressed comparably, with *PRICKLE2* being over-expressed in SCZ (p<0.05). *MCAM*, a DEX gene with lower expression in SCZ, encodes CD146, a strong WNT5A receptor for a different non-canonical signaling pathway that regulates cell migration (32). Over-expression of DEX gene protein kinase C could result from both Ca^2+^ and planar cell polarity (PCP) non-canonical Wnt signaling pathways.

### WGCNA analysis

WGCNA identified 23 gene expression modules containing 78.0% of expressed genes (18,636 genes). Module size ranged from 3,934 to 64 genes. Only module #3, containing 32 DEX genes (out of 2,675 genes in the module), showed significant enrichment of DEX genes (Fisher’s exact test; OR=3.6; raw p=4.0×10^−11^). This enrichment was driven by genes that were in the main cluster of DEX genes seen in the hierarchical clustering (Figure 2), including *WNT5A, GAS1*, and *FOXO3*. Based on GSEA, this module shows extremely significant enrichment for GO terms “cell cycle process”, “chromosome segregation”, and “DNA replication” (corrected p<5×10^−18^ for all), among others. Additionally, the module eigengene showed a significant correlation with SCZ status (p=0.004).

Hierarchical clustering of samples based on the expression of genes in WGCNA module #3 shows a clear separation into two groups, an effect further supported by examination of the heatmap (Figure 4A). The same separation is apparent when SCZ and CTL samples are examined separately (Figure 4BC). The smaller subgroup shows an enrichment of SCZ samples as compared to controls (Fisher’s exact test, p=3.6×10^−05^, OR=4.76).

**Figure 4.**
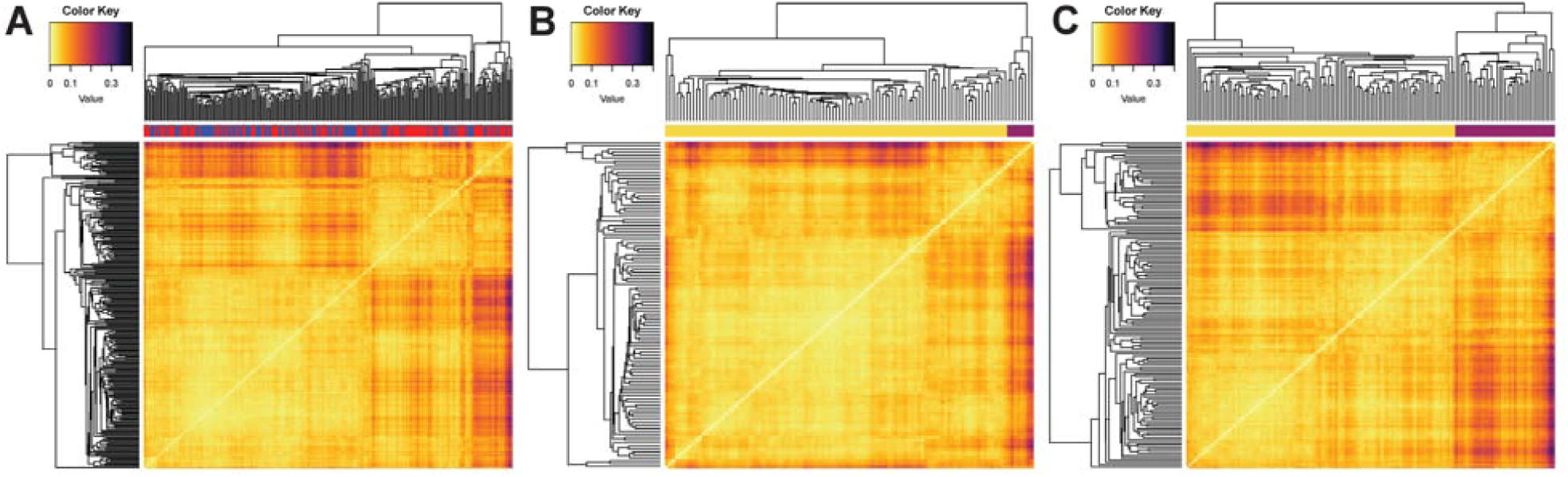
Heatmaps of sample-sample correlation based on genes from WGCNA module #3. **(A)** Heatmap for all samples. Color bar at top indicates SCZ (red) or CTL (blue) status. **(B)** Heatmap for only CTL samples. **(C)** Heatmap for only SCZ samples. Color bar at top indicates membership in a SCZ-enriched subset of samples (purple) or the main group of samples (yellow).

### Convergence with other genetic studies

Although SCZ GWAS is considered the most general and direct way to identify causative common genetic variants, consideration of other related phenotypes, and other types of genetic studies, such as *de novo* mutations, mutations segregating with psychiatric disorders in multigeneration families, and gene expression studies in relevant models also provide information about genes likely involved in SCZ. We found that many genes identified in these studies are also DEX in our study. *FOXO3, SRPK2, BACE2, GABRA4, RC3H1, NRXN3, TANC2* and *LYL1* are genome-wide significant in DEPICT-based association in GWAS meta-analysis of intelligence and associated traits (33). SNP within *PRKCA* is significant in GWAS of neuroticisms (34). Four DEX genes have been reported to have *de novo* non-silent mutations in individuals with SCZ (*PI15, TANC2* - (35), *ESAM, SSBP3 -* (36)), four have been reported to have *de novo* missense mutation(s) in patients with autism spectrum disorder (*MCAM, PRKCA* - (37), *FLG* (38), *PCDHB16* (39)), copy number variations of *IL1RAPL1* have been identified in multiple patients with SCZ (40) and multiple mutations in the same gene have been observed in individuals with autism spectrum disorders and intellectual disability (reviewed in (41)). An exonic deletion in *NRXN3* was found to segregate with neurodevelopmental and neuropsychiatric conditions in a three-generation Chinese family (42). Moreover, among 19 genes most correlated with WNT5A (R>0.6) nine have *de novo* mutations in either SCZ, autism spectrum disorder or developmental disorder, demonstrating striking convergence between gene expression data and *de novo* mutation studies of SCZ on a pathway level. In total, 18 of the 80 genes overlap with previously published results from genome-wide studies of psychiatric disorders, and some of them have supported evidence from several independent studies (*ESAM, FOXO3, SRPK2, IL1RAPL1, TANC2, PRKCA, NRXN3*).

The largest transcriptome study of SCZ was performed by the CommonMind Consortium (CMC) using post-mortem adult dorsolateral prefrontal cortex (258 SCZ vs. 279 CTL) (43), which has a very different pattern of gene expression than CNON or the fetal brain (Supplemental Figure S3, and (18)). Only two genes showed significant differences after correcting for multiple comparisons in both studies (*FAM110C* and *CLEC3B*) and the differences were in opposite directions in both cases. However, correlation of test statistics for genes expressed in both studies was highly significant (r=0.174; p<2.2×10^−16^, n=14924 genes); this correlation increased to r=0.42 (p<2.2×10^−16^, n=453 genes) on the set of genes that were nominally significant in both studies.

We also compared our results with data from studies of iPSC-derived neural progenitor cells, which are most similar to the first trimester samples in BrainSpan (44). We found three DEX genes, all changed in the same direction, in common with a study based on microarray expression profiling of 4 SCZ and 4 CTL individuals (*ARHGAP26, NRXN3, LRRC61*) (44), and there was a significant positive correlation in z statistics testing for differential expression between SCZ and CTL groups for genes with publicly available data (Pearson’s r=0.117, p=0.025, n=367 genes). This work was extended using RNA-Seq and two additional controls (4SCZ/6CTL) (45) and shared 5 genes with our DEX list (*GAS1, WNT5A, BACE2, FRMD6, PRKCA*); all genes except *PRKCA*, had the same direction of effect. As before, the overall comparison of test statistics with our study was significant (r=0.15, p=4.26×10^−5^, n=724 genes). This latter study implicated Wnt signaling in schizophrenia, which agrees with our findings of involvement of WNT5A in etiology of the disease.

However, the largest SCZ study using iPSC-derived NPCs (10SCZ/9CTL) from the same group is significantly negative correlated in test statistics with our data for all genes (r=-0.08, p<2.2×10^−16^, n=15,862). The results of this latest study are also significantly negatively correlated with two previous studies from the same group (r=-0.33, p=7.87×10^−12^, n=412 with microarray study and r=-0.19, p=1.13×10^−6^, n=654 with RNA-seq study).

## Discussion

Schizophrenia is a complex genetic disorder that originates during fetal development, typically manifests symptoms in adolescence and early adulthood, and persists throughout adult life. While post-mortem brain transcriptome studies assess changes in gene expression of differentiated neuronal and glial cells in the adult brain, we developed a genetically unmodified cell-based system, CNON, to study the neurodevelopmental component of the disorder. These cells, developed from olfactory neuroepithelium, are neural progenitors, and express a transcriptome most similar to the mid-fetal period of the brain (Supplemental Figure S3), a time of increased risk for the development of schizophrenia (46,47). We examined differences in mean gene expression between SCZ and CTL groups, but other approaches for finding biologically important differences are also possible (48). We identified 80 DEX genes at FDR<10% and found an overrepresentation of genes annotated with gene ontology terms related to processes of cell proliferation, migration and differentiation, all fundamental aspects of neurodevelopment.

We looked for convergence of our transcriptome data analysis with results of other types of genomic or transcriptomic studies of SCZ or related psychiatric diseases that were performed on a comprehensive, genome-wide or transcriptome-wide basis. GWAS is considered the most general way to identify causative genomic loci associated with common variants. The main mechanistic explanation for involvement of GWAS loci in disease etiology is through altering gene expression (20,21). However, the specific causal SNPs are generally not known, and neither are the genes which they regulate, or the developmental stage or type of cells where the regulation is important for the development of the disease. Despite these complexities, our study shows a significant agreement between our DEX gene list and genome-wide significant loci in the PGC SCZ2 GWAS (odds-ratio=3.8, p=0.049). Other genomic approaches, such as identification of *de novo* non-synonymous mutations in psychiatric disorders, also provide independent evidence for involvement of some DEX genes with SCZ or neurodevelopment in general, suggesting that alterations in expression of some genes and changes in gene function could result in similar phenotype. Finally, a transcriptome study (45) of iPSC-derived NPCs, a similar cellular model of SCZ, agrees with our conclusion of involvement of Wnt signaling in SCZ etiology.

Analysis of the DEX genes provides insight into the neurodevelopmental processes altered in SCZ. These genes appear to function in a number of biological pathways and functions, with the largest group being involved in Wnt signaling in general, and WNT5A signaling in particular. Most DEX genes in this group (co-expressed with DEX gene *WNT5A*), as observed in both the hierarchical clustering of DEX genes (Figure 2) and the WGCNA analysis (Figure 4), have known functions in Wnt signaling. The Wnt signaling pathway was also found to be over-represented in the PANTHER pathway analysis.

Wnt signaling is one of the most versatile signaling mechanisms involved in regulation of different cellular and organismal functions, including development of organs and tissues, balance between cell proliferation and differentiation, cell migration, and stem and progenitor cell maintenance. These functions are critical for proper brain development; for example, a crucial role of Wnt signaling in the developing cerebellum has been demonstrated (49). Our finding of a perturbation in Wnt signaling is consistent with previous studies (reviewed in (50–52), (53)). As presented above, Brennand and colleagues found a gene expression signal of altered Wnt signaling, using both microarrays (6) and RNA-Seq (45). Genomic disruption of *Disc1 (disrupted in schizophrenia 1)*, which segregates with psychiatric disorders including schizophrenia, results in an increased level of canonical Wnt signaling in neural progenitor cells (54). Gene expression in blood cells shows alterations in Wnt signaling in SCZ and BD, and plasma level of dickkopf 1 and sclerostin, known inhibitors of Wnt signaling, are decreased in patients (53). These studies and our results agree that Wnt signaling is enhanced in SCZ.

In CNON lines, *WNT5A* has the highest expression of Wnt ligand genes (TPM=185.5) and is DEX (FDR=1.9%, SCZ>CTL). Transcription of *WNT5A* is regulated by a JAK-STAT3 signaling pathway, which includes DEX genes *JAK1* (FDR=3%, SCZ>CTL) and Leptin (*LEP*, FDR=9.9%, SCZ>CTL) (Figure 3A) (25). Additionally, *IL-32* (FDR=0.58%, SCZ>CTL) has been shown to increase the expression of IL-6, another JAK-STAT3 ligand (26), and may increase WNT5A through IL-6 or via STAT3. Lastly, *PI15* (FDR=0.059, SCZ>CTL), has been shown to induce *WNT5A* (55). We also observe two DEX genes in the downstream canonical signaling pathway of WNT5A; PGC1-alpha (FDR=4.5%, SCZ>CTL) and FOXO3 (FDR=3.5%, SCZ>CTL), the latter is a product of one of the three DEX genes that lies under a PGC SCZ2 GWAS peak (Figure 3B), providing convergence between the results of the two types of studies at the level of signaling pathways. WNT5a is also involved in multiple interrelated non-canonical Wnt signaling pathways. There are two DEX genes whose products are involved in non-canonical Wnt signaling: *MCAM* (FDR=4.5%, CTL>SCZ), encoding cell surface glycoprotein CD146, and *PRKCA* (FDR=5.9%, SCZ>CTL), encoding protein kinase C alpha (PKC). CD146 is a high affinity receptor for WNT5A (32) and in conjunction with PKC regulates localized membrane retraction, establishing directionality of locomotion (32,56).

The DEX gene with the highest correlation with *WNT5A* is *GAS1*, growth arrest-specific protein 1. Transcription of this gene is induced by several Wnt ligands (57). Among the functions of GAS1 is attenuation of SHH signaling (57), one of key pathways in neurodevelopment. GAS1 is known to regulate the proliferation of the external germinal layer and Bergmann glia, influencing the size of the cerebellum (58), which has been observed to be altered in SCZ (59).

Analysis of individuals by hierarchical clustering and heatmap analysis for WGCNA module #3, which showed a significant enrichment of DEX genes, identifies a group of individuals with abnormal expression of genes involved either in WNT5A regulation or downstream Wnt signaling. This subset is significantly enriched for individuals with SCZ and may represent a molecular subtype.

In summary, our results show that DEX analysis of CNON cells produces biologically meaningful results, demonstrates convergence with other genome- and transcriptome-wide studies of SCZ and related traits, and provides insight into specific mechanisms of developmental aspects of the disease. We also show that CNON are a good cellular model to study developmental aspects of brain disorders. Further studies using this model will improve the mechanistic view of SCZ etiology with finer detail. CNON cells are derived from living individuals, providing numerous opportunities for personalized medicine at a substantially lower cost than the development of iPSCs. Cell lines provide additional opportunities to test hypotheses using molecular tools such as CRISPR, siRNA and miRNA knock-down. These technologies can be combined with our ongoing epigenetic studies (60) and functional tests for proliferation, migration, cell adhesion, and to evaluate cellular phenotypes in 2D and 3D cultures.

## Acknowledgments

This study was supported by a NARSAD Young Investigator Award and a NIMH grant MH086874 to OVE and NIMH grant MH086873 to JAK. Previous version of this manuscript is available at BioRxiv (doi: https://doi.org/10.1101/209197)

