## Supplemental material for "Gene expression in patient-derived neural progenitors implicates WNT5A signaling in the etiology of schizophrenia"

### Mapping and assignment of reads to genes

Every set of sequencing reads with unique index in every channel of flow cell was quality controlled for overall complexity (total number of reads, average entropy, percentage of reads with low entropy, frequency of most common K-mers and monomers). Prior to mapping, we removed reads containing more than 50% of adapter sequences, monomers or other low entropy reads (metric entropy below 1%). The rest of reads from each individual channel were trimmed (if adapters constitute less than 50% of the read) and sequentially aligned to rRNA, mtDNA, the rest of human transcriptome (GenCode v22 gene models) and genome (GRCh38) using our custom RNA-Seq alignment pipeline, GT-FAR v12 (<https://genomics.isi.edu/gtfar>). Mapping quality was monitored by examining the distribution of reads with different number of substitutions to match the reference. Reads mapped to rRNA and mtDNA were excluded from following analysis. Reads generated from the same library, but ran in different channels or different flow cells were assessed for quality separately, and those that passed QC were united as a set of reads specific to the individual. We required every library ran in one channel to have at least 1 million reads after QC, with overall mapping rate at least 75% and reads aligned to the correct strand at >90%. We analyzed female-specific (*XIST*) and male-specific (*DDX3Y*, *USP9Y*, *KDM5D*) ubiquitously-expressed genes to confirm samples properly clustered with annotated sex. We also required that RNA-defined genotypes corresponded to DNA-defined genotypes, which were previously determined by microarray or whole genome sequencing. Additionally, we identified and removed outliers with regard to the alignment percentage to mtDNA, rRNA and gene models. We vigorously tested correlation of gene expression between libraries, RNA samples, individuals, as well as channels and flow cells. We assumed that correlation between sequencing data should generally decrease in the following order: the same library in the same flow cell, the same library in different flow cells, different libraries from the same RNA sample, libraries from different RNA samples, libraries from RNA samples purified from different biopsies of the same individual, and lastly libraries from different individuals. Most sequencing data fit the expected pattern, and outliers were excluded after thorough investigation of potential reasons for the deviation. Finally, we analyzed the relationship between number of detected genes and number of aligned reads; a few outliers from a best-fit relationship were excluded. Reads that aligned to the sense strand of a single gene model with minimal number of mismatches (not more than six) were assigned to the gene. The number of reads assigned to a gene was used as a proxy of gene expression in DEX gene analysis.

### Differential gene expression analysis

A critical component of differential expression (DEX) analysis in complex genetically heterogeneous diseases such as schizophrenia is adequate correction for technical elements that increase noise and add confounding factors which often have a stronger effect than the disease itself. Adding covariates, which are used for correction of specific or unknown confounding factors, may increase power if sufficient variation is taken into account, or reduce it, if the covariate does not explain a substantial amount of variation, and there is a risk of overfitting that may overwhelm the benefits of adjustment if too many are added.

Without adjustment for any other variables, diagnosis accounts for 0.8% of variation in the total data. Known covariates analyzed for inclusion in the analysis are sex, age, race/ethnicity, library batch, and sequencing batch (flowcell). Sex and age account for 2.4% and 1.7% of total variance in a model including only themselves, respectively. Given the fact that both sex and age are significantly imbalanced with respect to diagnosis (Supplementary Table 1), both were included in the model. The combined model of diagnosis, sex, and age explained 3.4% of variation.

Based on the equation for the Bayesian information criterion (BIC) (1), a single added parameter must explain at least an additional 2.1% of total variance to reduce the BIC, indicating a better fitting model. This was used as a guideline to determine inclusion of additional parameters to adjust for known batches, such as samples that were part of the same library creation batch or samples that were run on the same flowcell. Three library batches were found to explain more than 2.1% of variation (each explains from 2.7% to 11.6%) and were added to the model. Four flowcells were found to explain more than 2.1% of variation in the data, however, library batches and flowcells are highly correlated, and library batch explained variance better in fewer covariates. Samples from the same library batch that were run on different flowcells appeared the same in PCA, while samples from different library batches that were run on the same flowcell appeared different. To deal with technical effects we added 3 additional covariates for the library batches; the updated model explained 20.1% of variation in the data.

Control and case groups were also not balanced by racial/ethnic category (Supplementary Table 1; Chi-Square test p = 0.02). However, analysis of the first 10 principal components for the log-transformed normalized expression values showed no substantial general effect of racial/ethnic category on gene expression (test for difference in means between groups by one-way ANOVA, all FDR > 10%), a drastic difference with the effect in genotyping data for a subset of samples (Supplemental Figure S7). The same was true when testing on the top 10 principal components after correction for library batch effects.

Differential expression analysis with racial/ethnic category included as a covariate (4 groups: Non-Hispanic Caucasian, Hispanic, African-American and Other) identified genes showing expression differences correlated with race/ethnicity, however the additional residual variance explained by the added covariates (1.2% of variance explained by 3 indicator variables) was insufficient to offset the penalty for increased parameterization in the model. All DEX genes identified using racial/ethnic category as a covariate showed differences in the same direction as DEX genes identified without correction for ethnicity (Pearson correlation of test statistics, r = 0.990) but generally of less statistical significance: 53 of 80 DEX genes remain at FDR < 0.1, 79 of 80 at FDR < 0.2 (Supplemental Figure S8). 24 additional genes became significant at FDR < 0.1 with the addition of racial/ethnic category covariates, also all showing trends of differential expression in the same direction as in the main analysis (76 of 77 at FDR < 0.2). Thus, overall influence of race/ethnicity on expression profile appears insignificant and does not justify correction for that covariate by racial ethnic category, which groups rare racial/ethnic backgrounds in a single "Other" category and does not account for the continuous distribution of racial/ethnic background such as that seen between Hispanic and non-Hispanic Caucasians (Supplemental Figure 6A). Furthermore, adding covariates for race/ethnicity results in only small differences in significance for all but 2 genes (Supplemental Figure S8), and test statistics are highly correlated both for all expressed genes (Pearson r = 0.972) and for DEX genes (Pearson r = 0.990). *SLPI* appears to have dropped greatly in significance due to the fact that, despite showing a decrease in SCZ in 3 of 4 racial/ethnic groups, it shows a slight increase in SCZ among Hispanics. *PADI2* appears to have greatly increased in significance because expression in Non-Hispanic Caucasians is significantly higher than in the other 3 groups.

To adjust for possible unknown confounders we used Surrogate Variable Analysis (2). We calculated how many SVA covariates we should use based on the method proposed by Leek (3) and implemented in the "num.sv" function of the SVA package while using the previous covariates as the preexisting model. The addition of one surrogate variable to previously included explicit covariates was recommended. Adding this calculated covariate to the model explained 31.4% of the total variation.

### Permutation analysis of differential expression

To assess the probability that our DEX findings could be due to random statistical variation, we performed two forms of permutation analysis: a standard permutation analysis in which some case/control imbalance between groups is expected by chance (and hence some effect of SCZ), and a second with comparisons where we can expect the null hypothesis to hold (case vs. case; control vs. control). For the first, we randomly permuted the diagnosis labels but held all other factors constant and found a median of 11 DEX genes at FDR < 10% (mean = 28.92; 25th percentile = 5; 75th percentile = 22). We used the Wilcoxon signed rank test to assess significance of the mean signed rank between permuted and experimental data, and the number of DEX genes found in permutation analysis is significantly lower (p < 2x10^-16^) than in the original analysis. This shows that the difference in gene expression between the actual CTL and SCZ groups is greater than between random groupings.

For the second permutation analysis, we compared groupings where the null hypothesis should hold (random half of SCZ vs another half SCZ and similar CTL-CTL; null comparisons) to those where we expect the null hypothesis to be violated (case/control comparisons). The null comparisons resulted in significantly fewer DEX genes at FDR < 10% (SCZ-SCZ, median=2.0, mean=2.7; CTL-CTL median=2.0, mean=3.7) as compared to case/control comparisons (medians = 2.2 and 4.4 DEX genes, means = 5.3 and 8.3, for two sample sizes corresponding to either half of controls or half of SCZ, respectively; p < 3.6e-15 by Wilcoxon test).

DEX genes from the main analysis were found to be differentially expressed (FDR < 10%) in case/control subsets permutations much more often than in null comparisons (median = 13.7% for case/control compared to median = 0.2% for null comparisons; Mann-Whitney test p < 2.2e-16). 33 DEX genes were not found to be significant in any null comparisons, while 2 (*CHI3L1* and *LINC01013*) were found to be significant in at least 10% of null comparisons. Comparatively, only 1 DEX gene (*FAM110C*) was not found to be significant in any case/control comparisons, while 62 were found to be significant in at least 10% of case/control comparisons.

These permutation analyses indicate that the majority of genes found to be DEX are not likely to be false positives and, instead, are reproducible results. However, the fact that the probability for DEX genes to be identified in case-control permutations is not high indicates that power is relatively low at that sample size, suggesting many more DEX genes exist.

| **Supplemental Table S1. Descriptive statistics for individuals in the study.** Balance between SCZ and control groups was tested by Chi-Square test (sex and race/ethnicity) or by Welch's t-test (age). | | | |
| --- | --- | --- | --- |
|  | **Controls** | **SCZ** | **Balance Test P-value** |
| **Total** | 111 | 144 |  |
| **Sex** |  |  | 0.025 |
| Male | 67 (60.4%) | 106 (73.6%) |  |
| Female | 44 (39.6%) | 38 (26.4%) |  |
| **Age** | 49.9 (S.D. = 12.7) | 40.5 (S.D. = 12.1) | 6.42E-09 |
| **Race/Ethnicity** |  |  | 0.02 |
| Non-Hispanic Caucasian | 44 | 41 |  |
| African-American | 27 | 62 |  |
| Hispanic | 27 | 27 |  |
| Other | 13 | 14 |  |

**Supplemental Table S2. Mean gene expression of marker genes in CNON cells.** TPM: transcripts per million.

| **Gene Symbol** | **Marker name** | **TPM** |
| --- | --- | --- |
| ***glial markers*** | | |
| S100B | S100 beta | 0.09 |
| GFAP | GFAP | 0.16 |
| OLIG2 | Olig2 | 0.17 |
| ***epithelial markers*** | | |
| KRT5 | cytokeratin-5 | 0.09 |
| CDH1 | CDH1 | 0.21 |
| ***Neuronal markers*** | | |
| UCHL1 | PGP9.5 | 173.55 |
| MAP1A | MAP-1a | 26.22 |
| MAP1B | MAP-1b, MAP5 | 125.5 |
| TUBB3 | β-tubulin 3 | 2.35 |
| CDH2 | N Cadherin | 56.48 |
| ***Markers of differentiated neurons*** | | |
| OMP | OMP | 0.5 |
| GNAL | Golf alpha | 0.88 |
| RBFOX3 | NeuN | 0.31 |
| TH | Tyrosine hydroxylase | 0.28 |
| ***Neural progenitor markers*** | | |
| NES | Nestin | 61.21 |
| VIM | vimentin | 3777.85 |
| REST | REST | 44.86 |
| NEPRO | NEPRO | 26.67 |
| ***Notch signaling*** | | |
| NOTCH2 | Notch2 | 70.21 |
| PSEN1 | Presenilin 1 | 25.53 |
| PSEN2 | Presenilin 2 | 14.23 |
| ADAM10 | ADAM10 | 44.66 |
| ADAM17 | ADAM17 | 39.94 |
| JAG1 | Jagged1 | 11.91 |
| ***Cell proliferation markers*** | | |
| MKI67 | Ki-67 | 75.98 |
| CCND1 | Cyclin D1 | 537.46 |
| CCNB1 | Cyclin B1 | 288.61 |
| ***Neural Precursor Cell Expressed, Developmentally Down-Regulated (NEDD) ubiquitin protein ligases*** | | |
| NEDD4 | NEDD4 | 56.05 |
| NEDD9 | HEF1 | 13.85 |
| NEDD8 | NEDD8 | 116.96 |
| NEDD1 | NEDD1 | 59.77 |
| NEDD4L | NEDD4-2 | 13.65 |
| ***Stemness and proneural markers*** | | |
| POU5F1 | Oct-4 | 2.19 |
| SOX2 | SOX2 | 0.38 |
| ASCL1 | Mash1 | 0.21 |
| NANOG | Nanog | 0.35 |
| ATOH1 | MATH1 | 0.18 |
| NEUROD6 | MATH2 | 0.19 |
| NEUROD4 | Neuro D4 | 0.15 |
| ATOH7 | MATH5 | 0.31 |
| NEUROG1 | Neurogenin 1 | 0.24 |
| NEUROG2 | Neurogenin 2 | 0.17 |

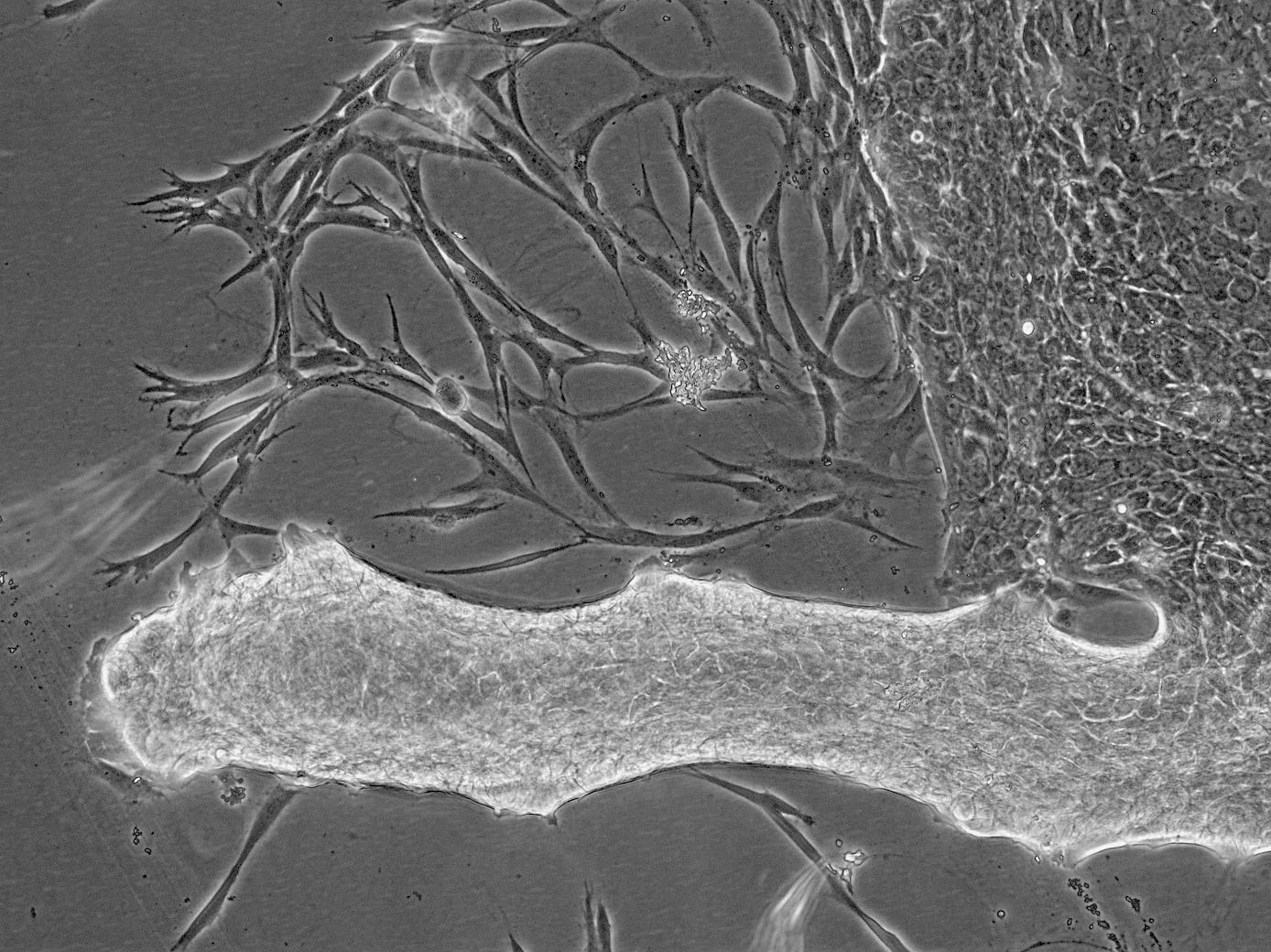

**Supplemental Figure S1. Neural progenitors** (white arrow) **and epithelial cells** (blue arrow) **growing out of biopsy piece** (red arrow).

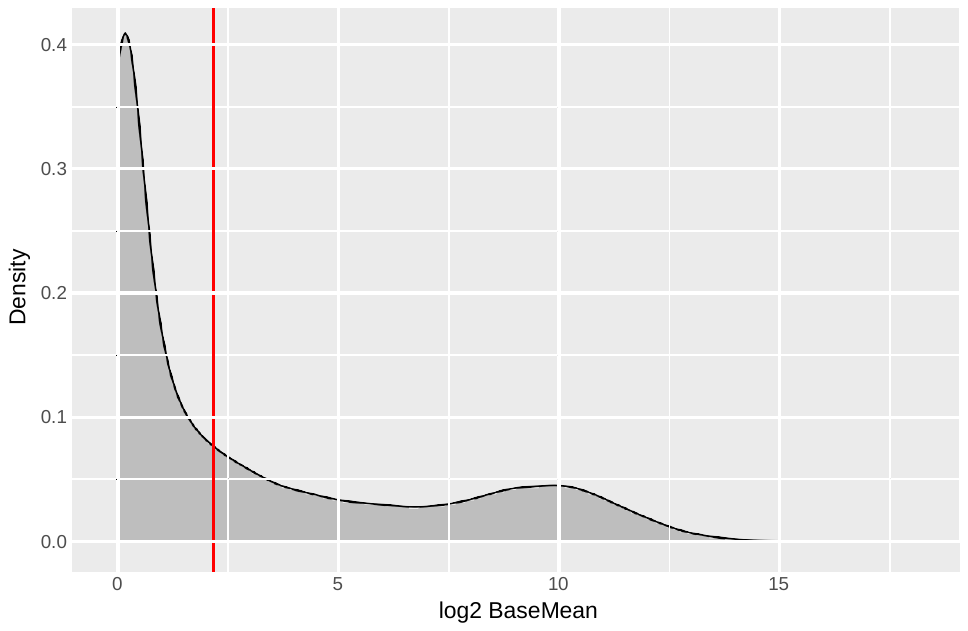
 **Supplemental Figure S2. Density plot of mean gene expression per gene.** Red line indicates the gene expression cutoff of 3.5 counts per sample on average (baseMean), which is equivalent to 0.17 counts per million (CPM). Cutoff was chosen liberally to include even low-expressed genes while still removing the large peak of unexpressed genes.

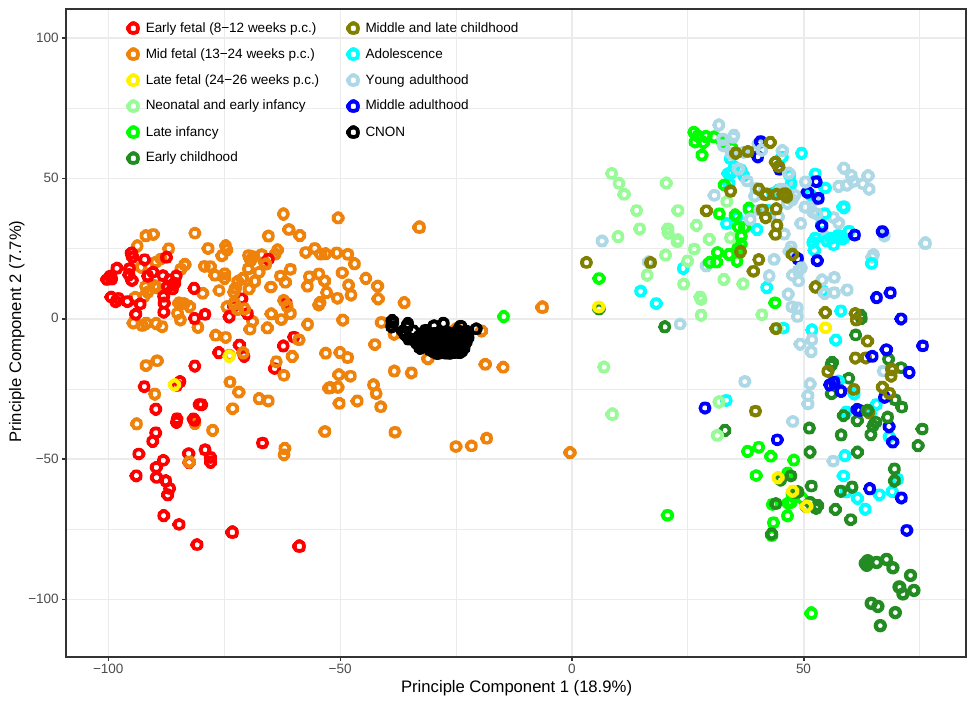
 **Supplemental Figure S3. Projection of CNON gene expression profiles (black) onto the first two principle components of BrainSpan data (colored).** Color represents developmental stages. Weeks p.c. is weeks post conception. Figure shows CNON samples cluster with mid-fetal brain samples (orange).

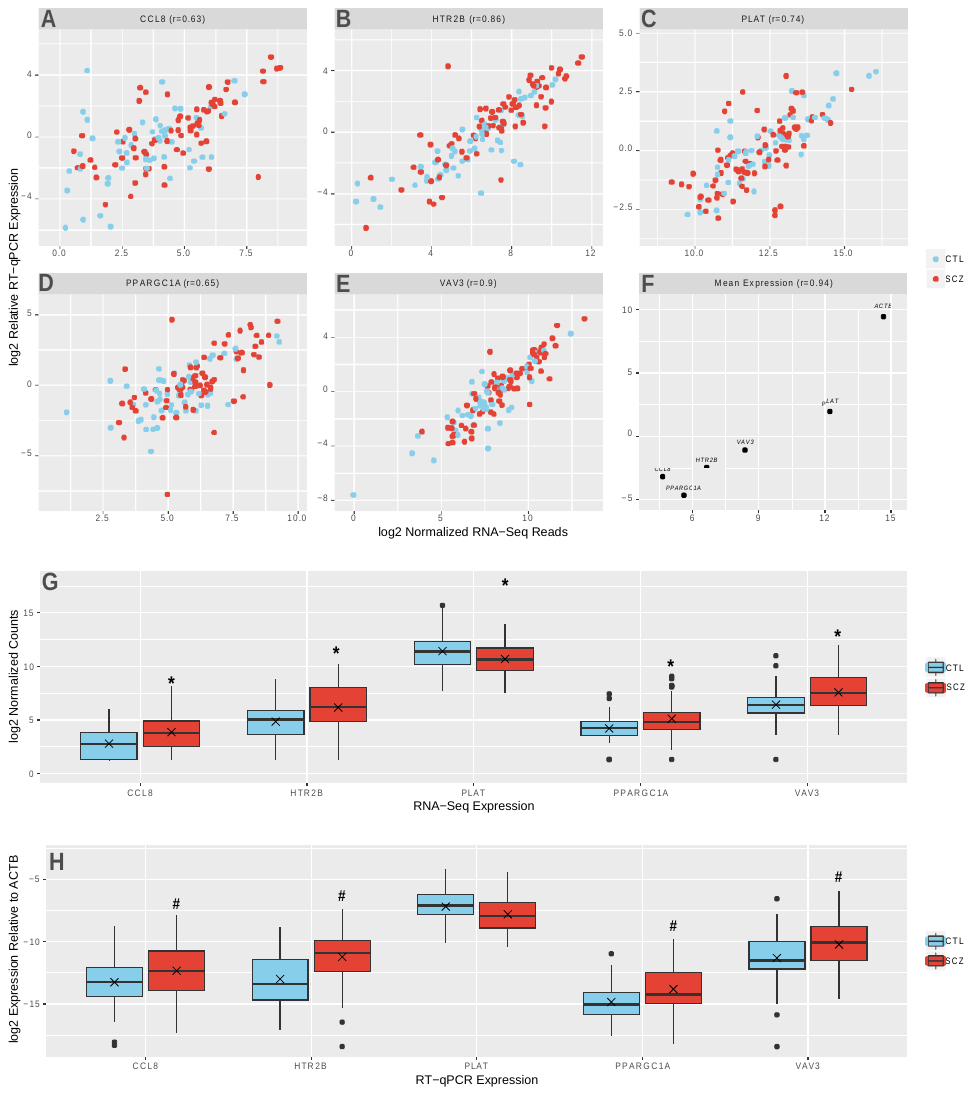
 **Supplemental Figure S4. RNA-Seq and RT-qPCR data of *CCL8*, *HTR2B*, *PLAT*, *PPARGC1A* and *VAV3* in the same 146 RNA samples normalized to expression level of *ACTB*. (A-E)** scatterplots showing correlation between expression measurements in RNA-Seq and RT-qPCR on samples. **(F)** scatterplot showing correlation of log-transformed average expression in RNA-Seq to deltaCt per gene. "r=" in header indicates Pearson correlation coefficient. **(G-H)** boxplots showing expression in samples divided into control (CTL) and schizophrenia (SCZ) groups. "x" indicates mean expression. * designates statistically significant difference in mean gene expression in RNA-Seq data (FDR < 10%), and # designates significance of mean gene expression measured by RT-qPCR (p < 0.05).

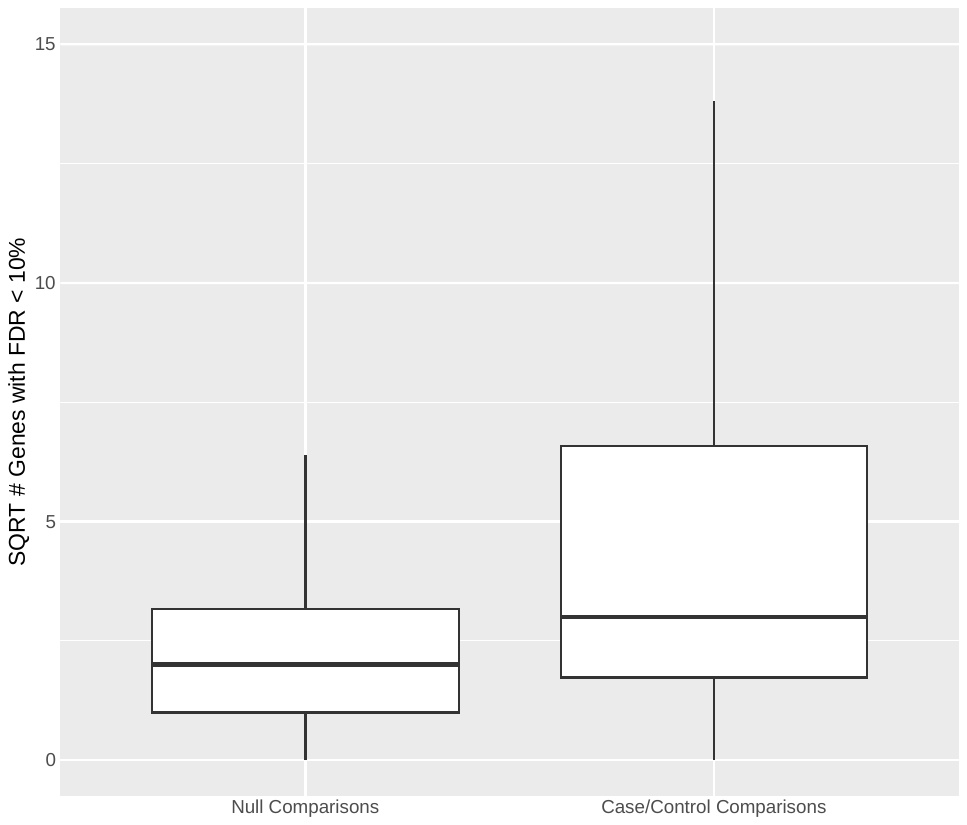
 **Supplemental Figure S5. Boxplot of the number of genes found to be DEX in subsets permutation analysis.** There were significantly more genes found DEX in comparisons between subsets of SCZ samples and subsets of CTL samples (case/control comparisons, right; SCZ-SCZ, median=2.0, mean=2.7; CTL-CTL median=2.0, mean=3.7) vs. comparisons between subsets of SCZ samples or subsets of CTL samples (null comparisons, left; medians = 2.2 and 4.4 DEX genes, means = 5.3 and 8.3, for two sample sizes corresponding to either half of controls or half of SCZ) (Mann-Whitney test, p < 3.6x10^-15^). Data points more than 1.5 inter-quartile ranges from the median are not shown.

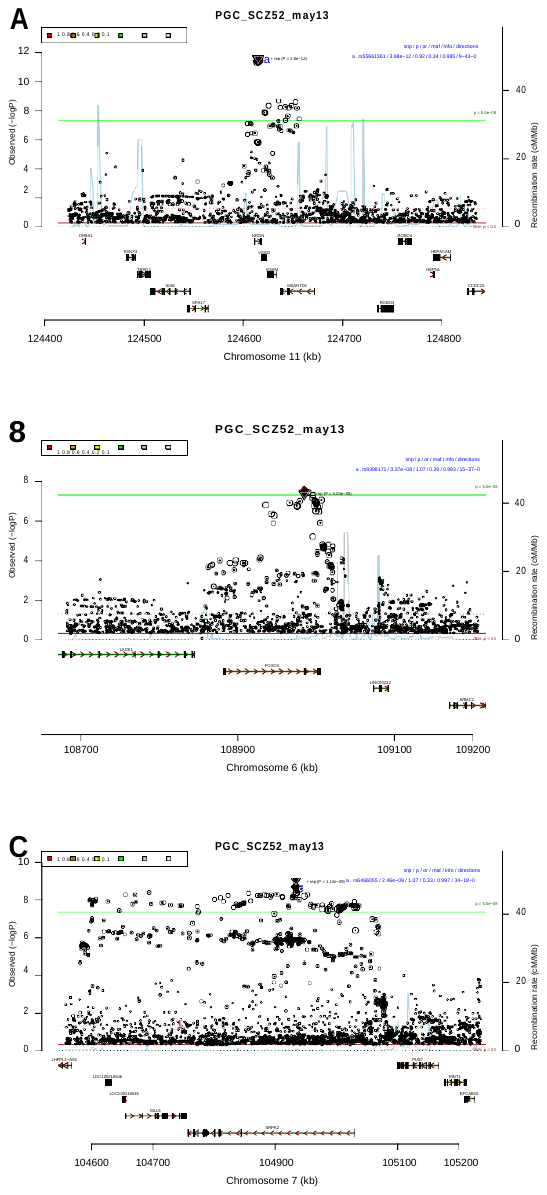

**Supplemental Figure S6. PGC SCZ2 GWAS region plots of DEX genes overlapping genome-wide significant variants.** **(A)** *ESAM* shows decreased expression in SCZ (FDR = 3.79x10^-6^). **(B)** *FOXO3* shows increased expression in SCZ (FDR = 3.49%). **(C)** *SRPK2* shows increased expression in SCZ (FDR = 4.91%). Figures generated by Ricopili from the Broad Institute (<https://data.broadinstitute.org/mpg/ricopili/>).

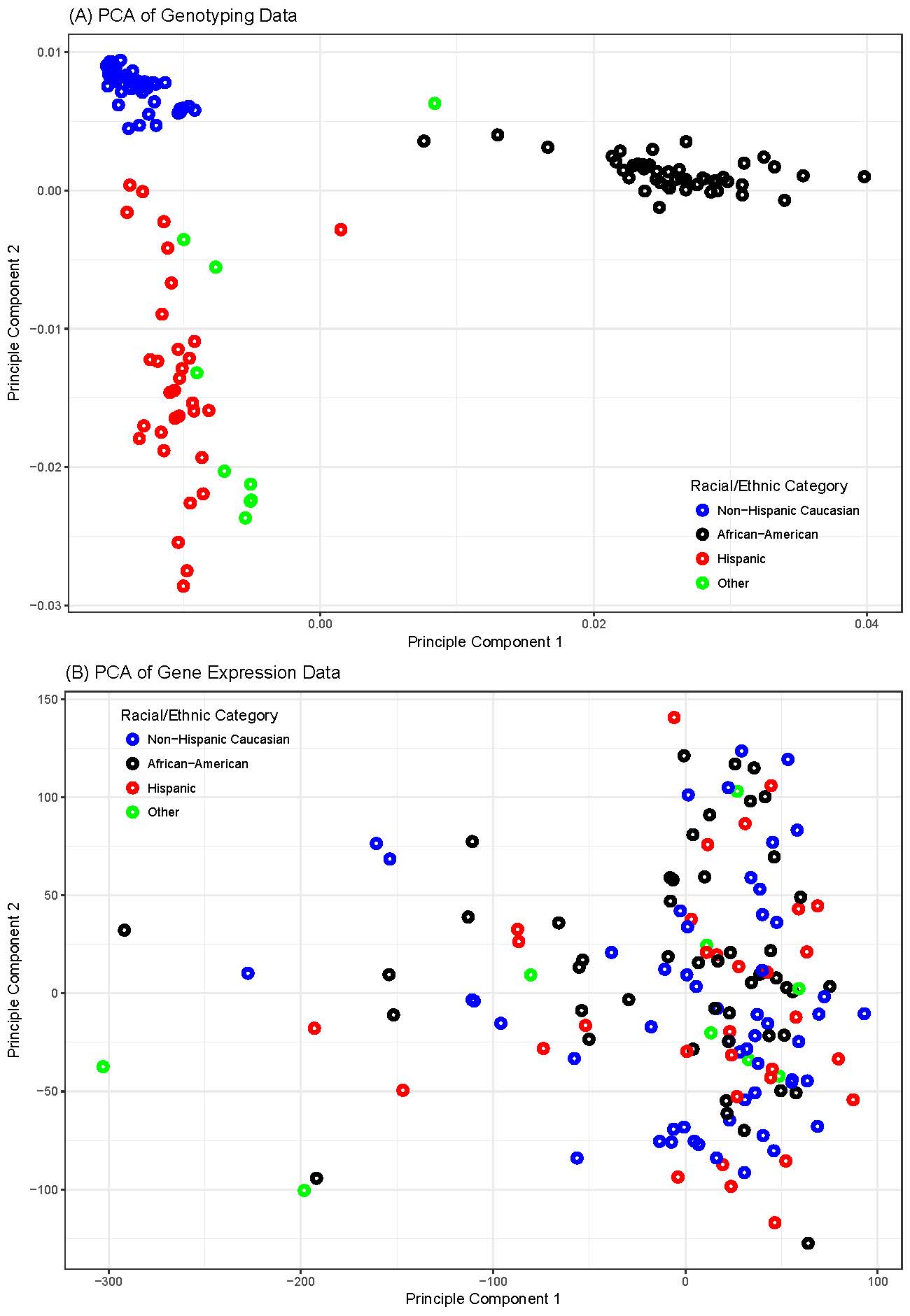

**Supplemental Figure S7. PCA plots of genotyping and gene expression data colored by racial/ethnic category. (A)** PCA of genotyping data. **(B)** PCA of gene expression data. Points are independent samples.

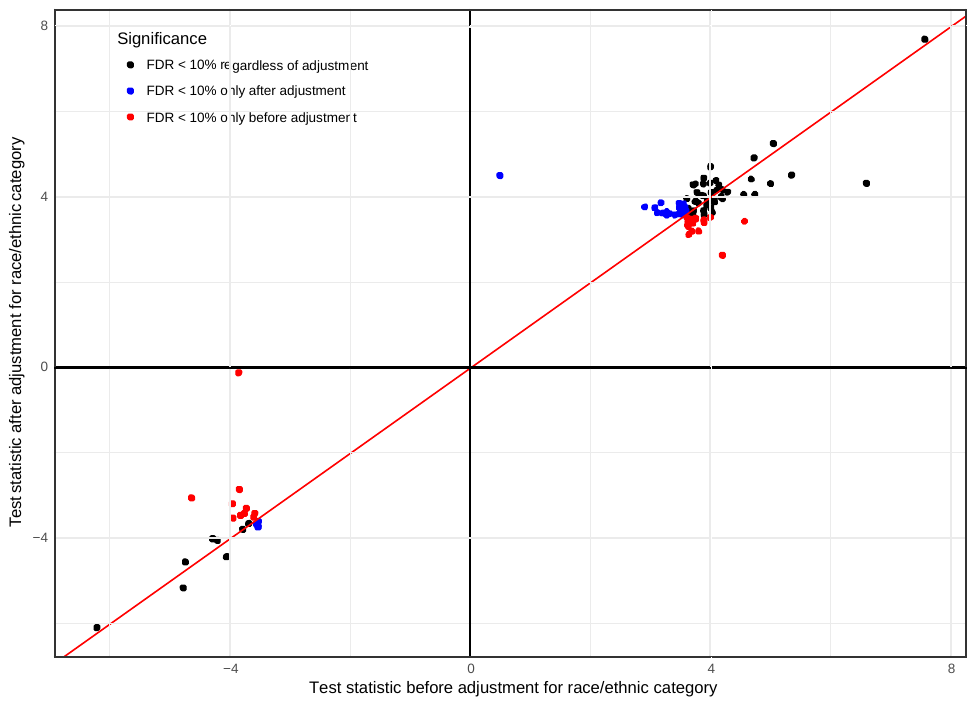

**Supplemental Figure S8. Scatterplot of test statistics (z-statistics) comparing results in analyses with and without including covariates for racial/ethnic category.** Black dots indicate genes that were significant (FDR < 10%) whether or not such covariates were used (53 genes). Red dots indicate genes that dropped out of significance with the addition of the covariates (27 genes). Blue dots indicate genes that entered into significance with the addition of the covariates (24 genes).
